## Supplementary Figures for "HERV activation segregates ME/CFS from fibromyalgia while defining a novel nosologic entity"

### Supplementary Materials

#### HERV activation segregates ME/CFS from fibromyalgia and defines a novel nosological entity for patients fulfilling both clinical criteria

Karen Giménez-Orenga[1], Eva Martín-Martínez[2], Lubov Nathanson[3], Elisa Oltra[4]*

[1]Escuela de Doctorado, Universidad Católica de Valencia San Vicente Mártir, Valencia, Spain.

[2]National Health Service, Manises Hospital, Valencia, Spain.

[3]Institute for Neuro-Immune Medicine, Dr. Kiran C. Patel College of Osteopathic Medicine, Nova Southeastern University, Fort Lauderdale, FL 33328, USA.

[4]Department of Pathology, School of Health Sciences, Universidad Católica de Valencia San Vicente Mártir, Valencia, Spain.

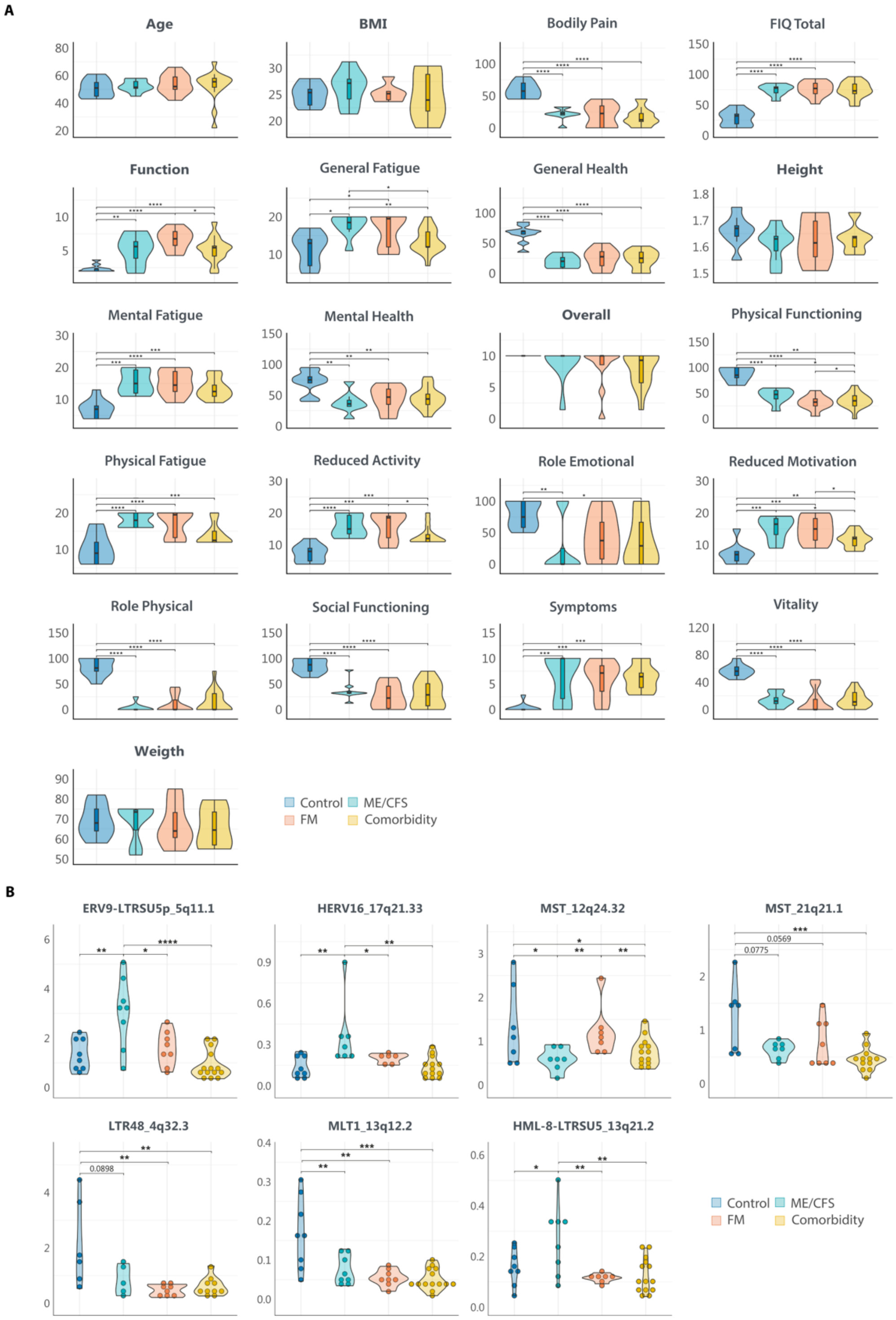


**Figure S1. Differences between patients and controls. (A)** Clinical data for participating subjects. Violin plots summarize the distribution for demographic variables (age, BMI, and weight), FIQ subdomains (Function, Overall and Symptoms) or total FIQ; for MFI subdomains (GF: general fatigue, Pfa: physical fatigue, RA: reduced activity, RM: reduced motivation, and MF: mental fatigue), and for SF-36 subdomains (PF: physical functioning, RP: role physical, BP: bodily pain, GH: general health, VT: vitality, SF: social functioning, RE: role emotional, and MH: mental health). Variables are ordered alphabetically. **(B)** Validation of DE HERV loci by RT-qPCR. Violin plots summarize the distribution and expression level of ΔΔCt values for each sample in each study group. Statistical tests: unpaired two-sample Wilcoxon Test with Benjamin-Hochberg p-value correction. (*$p<0.05$, **$p<0.01$, ***$p<0.001$, ****$p < 0.0001$).

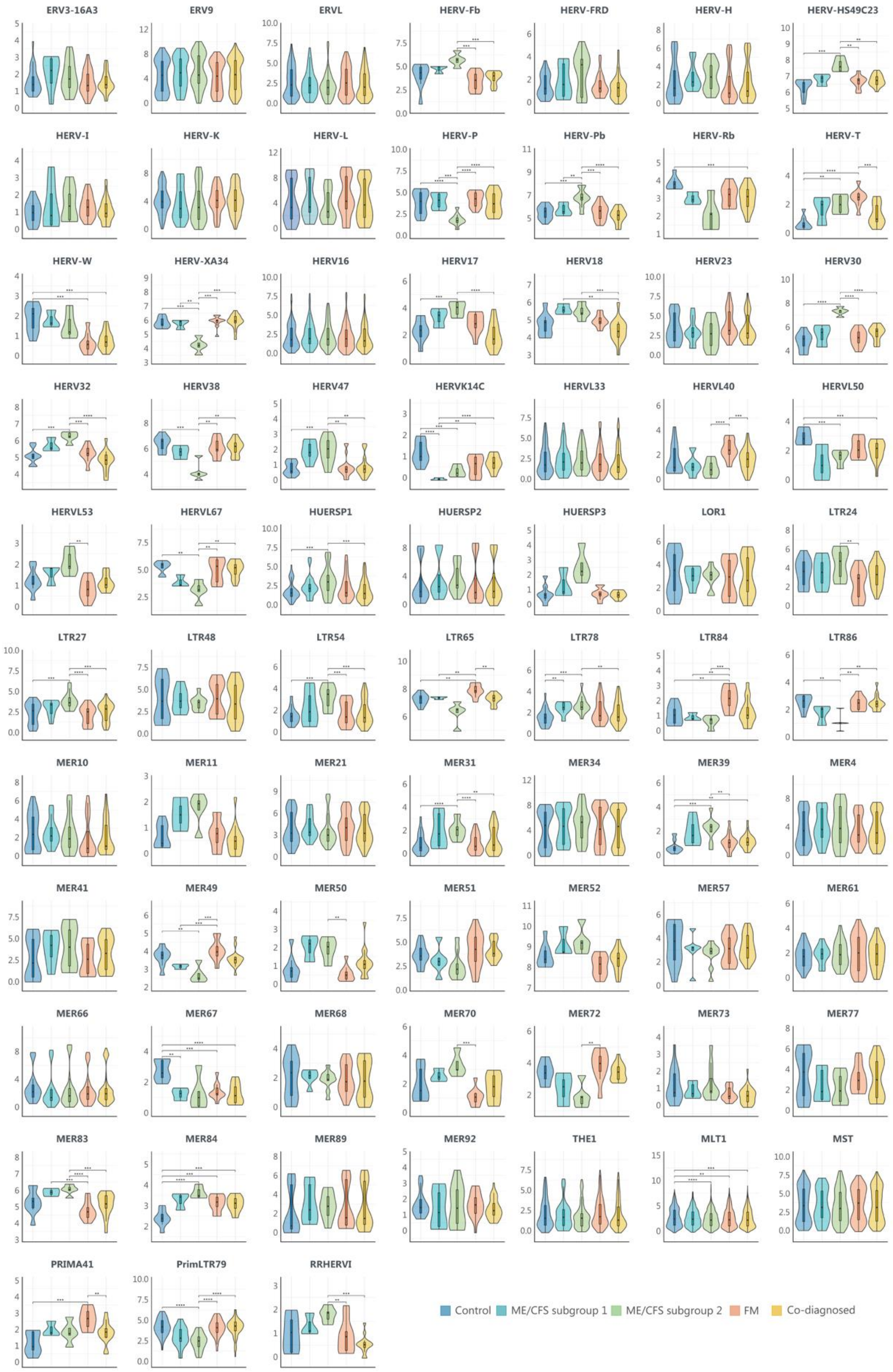


**Figure S2. DE HERV families by study group.** Violin plots summarize the distribution and expression levels of HERV probesets ($log_2$(mRNA expression level)) from a specific family for each sample in each study group. Statistical tests: unpaired two-sample Wilcoxon Test with Benjamin-Hochberg p-value correction. (*p<0.05, **p<0.01,***p<0.001, ****p < 0.0001).

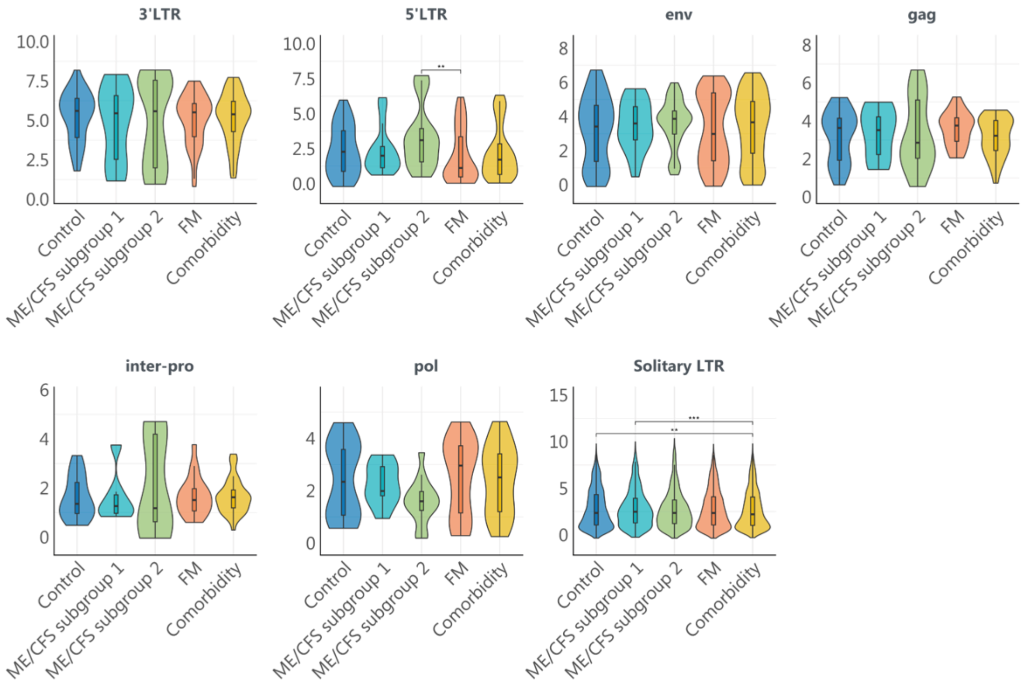


**Figure S3. DE HERV structures by study group.** Violin plots summarize the distribution and expression levels of HERV probesets ($log_2$(mRNA expression level)) from a specific HERV structure for each sample in each study group. Statistical tests: unpaired two-sample Wilcoxon Test with Benjamin-Hochberg p-value correction. (*p<0.05, **p<0.01, ***p<0.001, ****p < 0.0001).

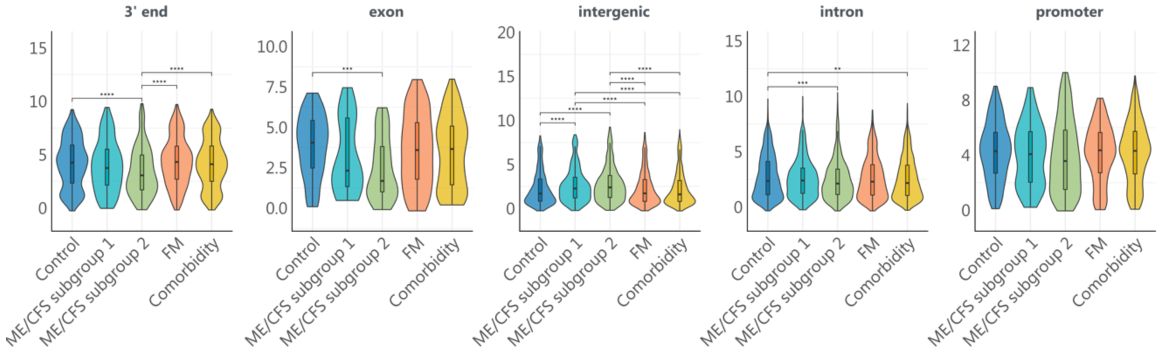


**Figure S4. DE HERV genomic location by study group.** Violin plots summarize the distribution and expression levels of HERV probesets ($log_2$(mRNA expression level)) from a specific HERV structure for each sample in each study group. Statistical tests: unpaired two-sample Wilcoxon Test with Benjamin-Hochberg p-value correction. (*p<0.05, **p<0.01, ***p<0.001, ****p < 0.0001).

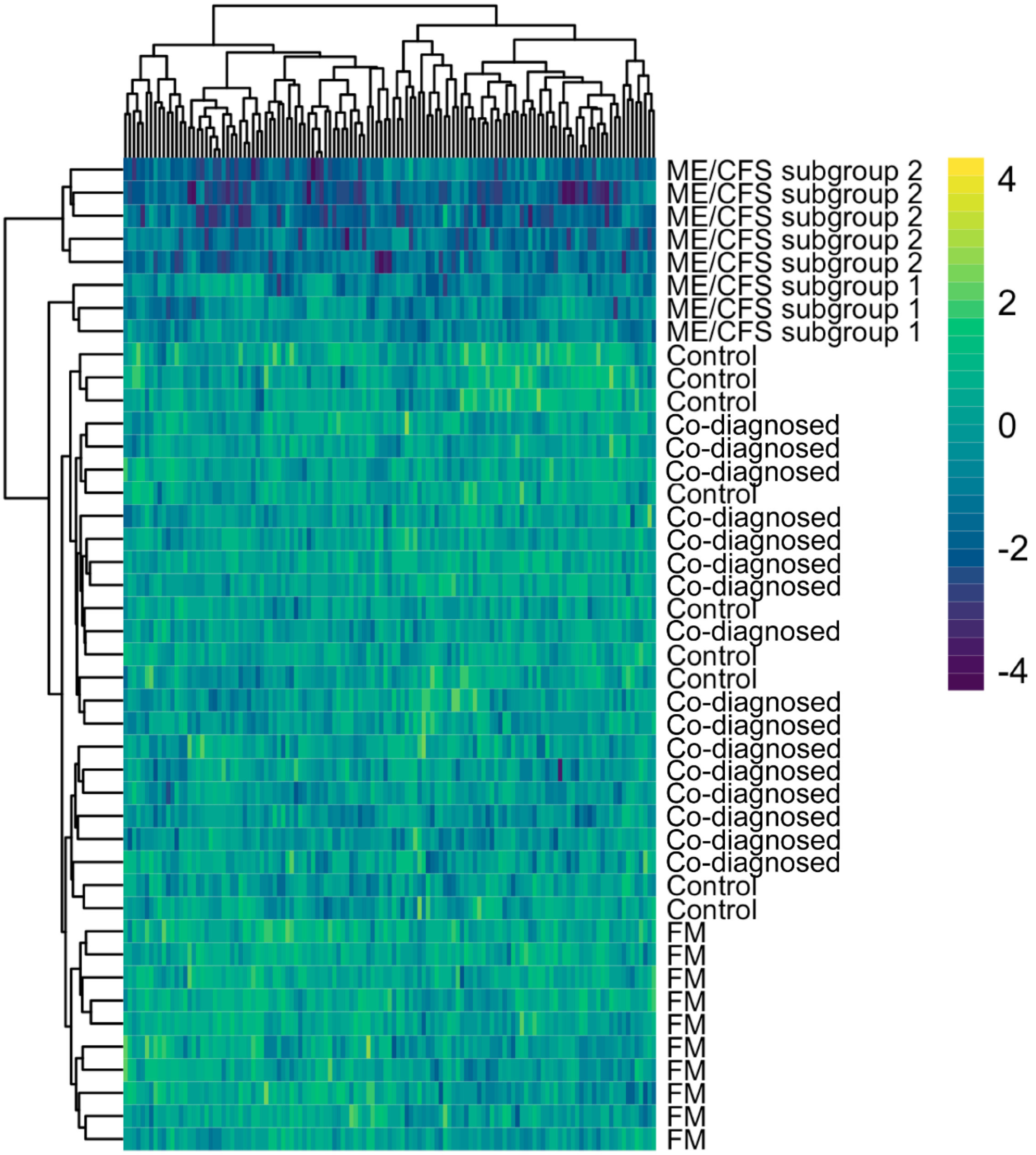


**Figure S5.** Intergenic solo LTR DE across study groups. Intergenic solo LTR expression heatmap of ME/CFS (n=8, green), FM (n=10, orange), co-diagnosed (n=16, yellow), and healthy control (n=9, blue) samples. The heatmap includes all solo LTR probes displaying significant DE between at least two of the compared groups (FDR<0.1 and |log2FC|>1).

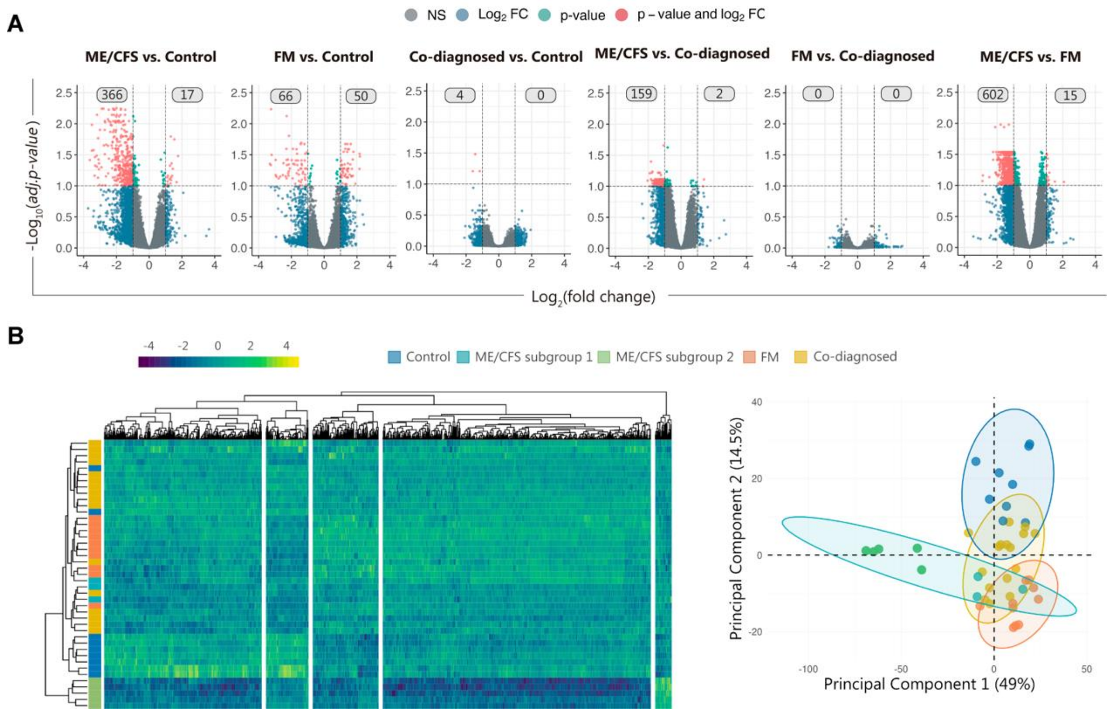


**Figure S6. Differential gene expression for ME/CFS, FM, comorbidity groups. (A)** Volcano plots showing the Log2(fold change) and the adjusted p-value for all the genes assessed by HERV-V3 microarray in each comparison by condition. Red dots indicate differentially expressed genes (FDR<0.1 and |log2FC|>1). Grey boxes indicate the number of upregulated and downregulated HERVs. (B) Gene expression heatmap and principal component analysis of ME/CFS (n=8), FM (n=10), comorbidity (n=16), and healthy control (n=9) samples. Sample groups are color-coded in green, orange, yellow, and blue, respectively. The heatmap and PCA include all gene probes displaying significant differential expression between at least two groups (FDR<0.1 and |log2FC|>1).

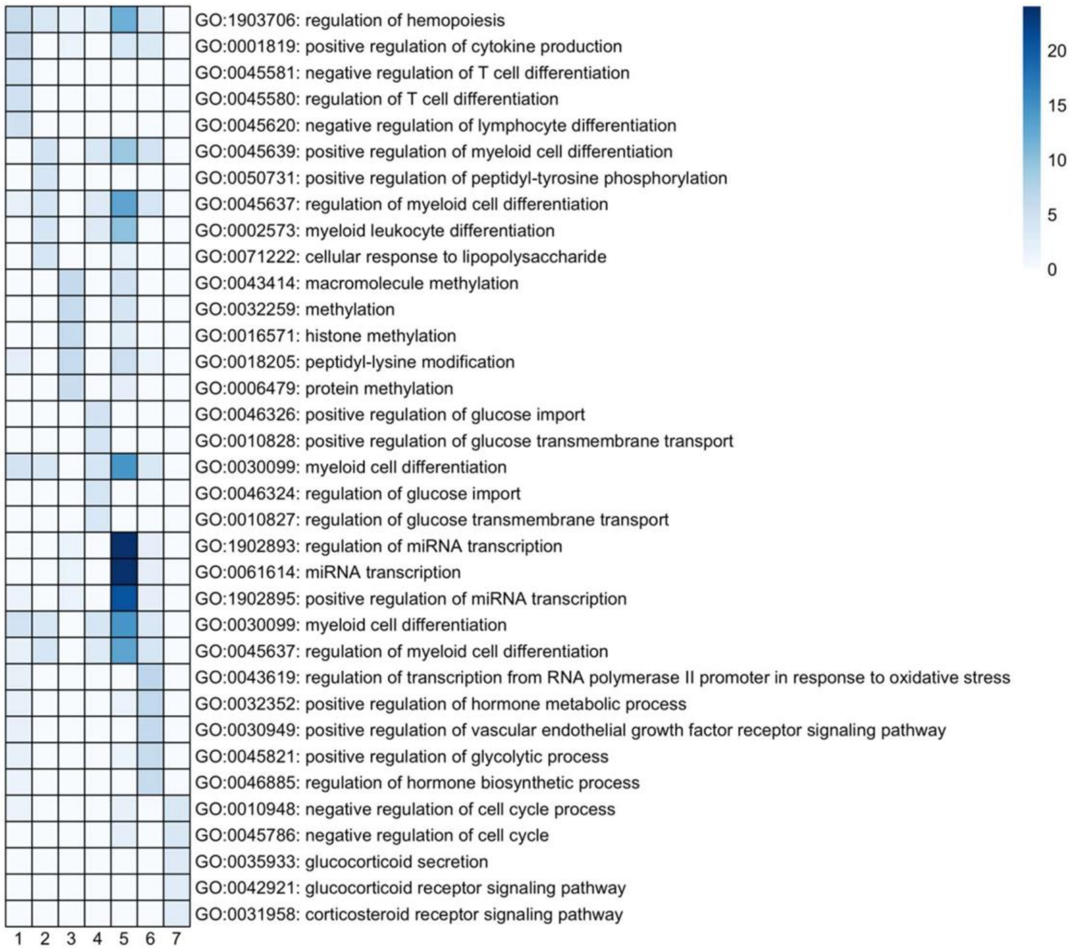


**Figure S7.** Gene ontology analysis of transcription factors binding sites enriched in HERV loci of modules one to eight.
